# The coordination of cell proliferation and cell-division orientation controls *Arabidopsis* radial style development

**DOI:** 10.1101/2024.12.10.627760

**Authors:** Iqra Jamil, Samuel W.H. Koh, Jitender Cheema, Laila Moubayidin

## Abstract

The biological mechanisms responsible for correct shape acquisition at the apex of the female reproductive organ—the gynoecium— remain poorly understood, despite its fundamental importance for successful plant reproduction and seed production. This process involves a rare bilateral-to-radial symmetry transition in *Arabidopsis thaliana,* orchestrated in part by the transcription factor SPATULA (SPT). Here, we show that SPT negatively controls cell proliferation, promoted by the hormone cytokinin, to enhance the robustness of cell-division orientation by orchestrating a coherent feed-forward loop that converges on the cell-cycle regulators CYCLIN-P3;1 (CYCP3;1) and CYCP3;2. While cytokinin induces both P-type cyclins, SPT represses their expression. Overexpression of CYCP3s disrupts style radial symmetry, causing the split-style phenotype and hypersensitivity to cytokinin observed in the *spt* mutant. Finally, we demonstrate a genetic link connecting the machinery of cell-division orientation, controlled by auxin, with the cell-proliferation input induced by cytokinin. Thus, our work reveals how the antagonistic auxin-cytokinin interaction scales up symmetry from the cellular to the organ level.

**Teaser:** Radial shape acquisition at the top of the plant female reproductive organ requires repression of CYCLIN-P3;1 and CYCLIN-P3;2.

## MAIN TEXT

### Introduction

Establishing the appropriate symmetry types, *e.g*., radial, biradial or bilateral symmetry, alongside specification of the polarity axes and tissue proliferation poses a significant challenge during the morphogenesis of plant and animal organs. In *Arabidopsis thaliana*, the bHLH transcription factor SPATULA (SPT)(*1*) plays a crucial role in promoting the development of the apical part of the female reproductive organ, the gynoecium, by supporting patterning and tissue specification along the medial and adaxial body-axis directions to form the radially symmetric style(*2–4*).

The development of the cylindrical style structure (Fig. 1A) requires dynamic control of auxin distribution via biosynthesis(*5, 6*),signalling(*7, 8*), and transport(*2*). SPT functions are instrumental during the apical fusion of the two carpels via directing auxin accumulation at the gynoecium apex(*2, 3*). We recently showed that the SPT-mediated auxin accumulation at the medial-apical cells is required to maintain orientation of cell division in the periclinal direction, *i.e.*, perpendicular to the apical-basal direction of organ growth(*9*). This is in line with the documented anisotropic growth at the style(*10, 11*). Growth analysis studies of the *Arabidopsis* style highlighted that growth is largely orchestrated along the apical-basal body axis. Accordingly, two upstream regulators of SPT, the *O*-glycosyl transferases SECRET AGENT (SEC) and SPINDLY (SPY) have been recently shown to promote style elongation and cellular expansion(*12*), consistent with a role for SPT in local adjustment of cell-division and growth.

**Fig. 1.**
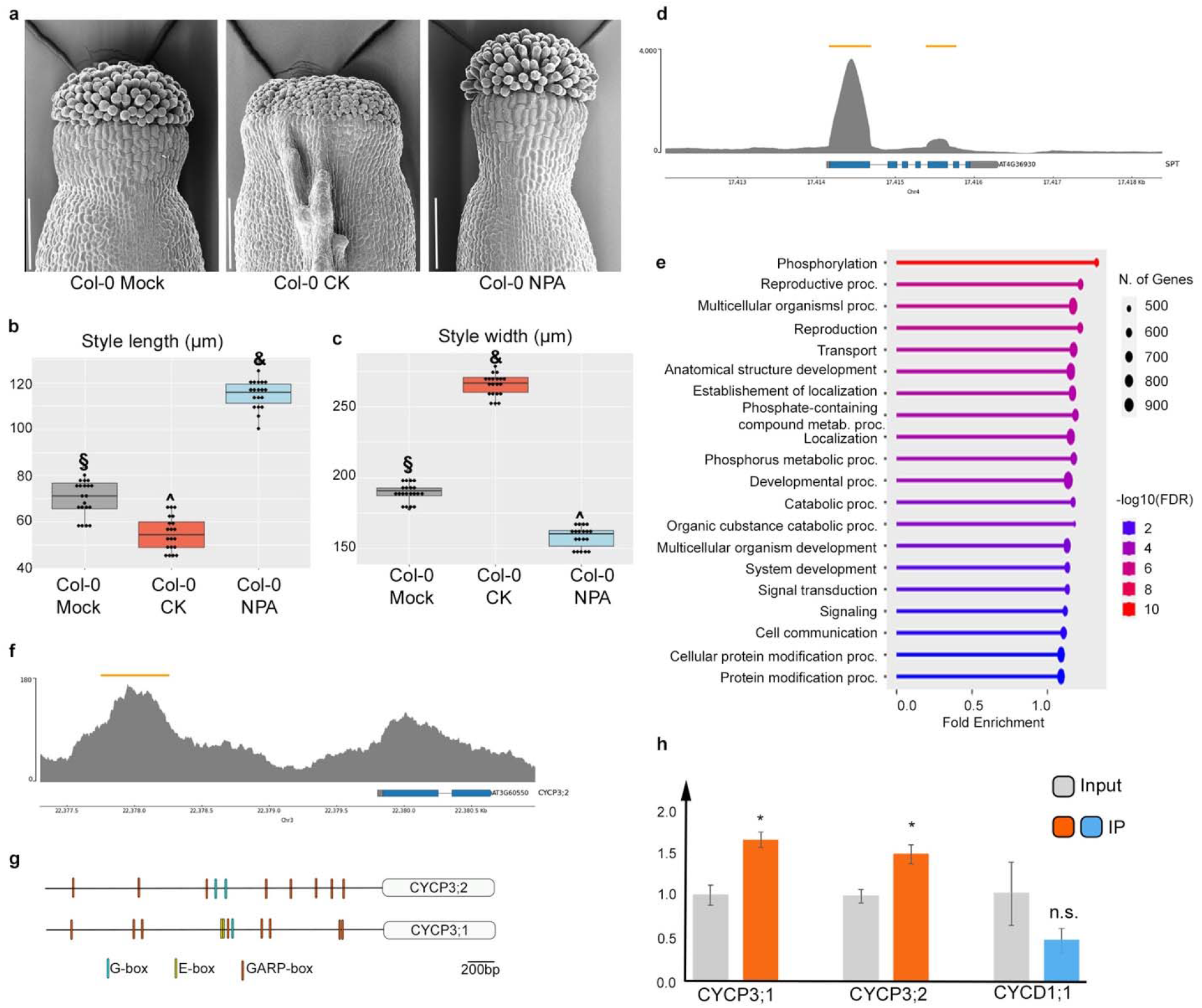
SPATULA direct targets in *Arabidopsis* inflorescences include *CYCP3;1* and ***CYCP3;2*** **(A)** Scanning electron microscope (SEM) images of wild-type Col-0 styles treated with mock (left panel), CK (middle panel, 50 μM BAP), and NPA (right panel, 100 μM). Scale bars represent 100 μm. (**B, C)** Boxplots of quantification of style length (b) and width (c) of samples depicted in (a). n=3 biological replicates; results of one representative biological replicate are plotted. Dots on boxes represent number of samples analysed per treatment. Symbols (&,§,^) on boxplots represent results of one-way ANOVA followed by Tukey’s Honestly significance difference (HSD) test. Tukey’s HSD *p*-values for Col-0 mock vs Col-0 CK, Col-0 mock vs Col-0 NPA, and Col-0 CK vs Col-0 NPA for both style length and width are <0.001 (&,§,^). (**D, F**) Representative raw Chromatin Immunoprecipitation sequencing (ChIP-seq) peaks of control gene *SPATULA* (*SPT*) and an *Arabidopsis* P-type cyclin gene i.e., *CYCLIN-P3;2* (*CYCP3;2*). n=3 biological replicates; peaks of one representative replicate are shown. Yellow bars on top represent peaks position on chromosome. Blue bars on bottom show exons, and grey lines show introns. **(E)** Bar chart of Gene Ontology (GO) terms of biological processes enriched in the set of 6,731 presumptive direct targets of SPT. ShinyGO (v0.80) was used to perform GO terms with the FDR cutoff 0.05. **(G)** Schematic representation of conserved motifs in the 5’ promoter regions of *CYCP3;1* and *CYCP3;2*. Different coloured bars on promoters represent binding sites for bHLH (G-box in cyan and E-box in yellow) and ARR-Bs transcription factors (GARP-box in orange). Note the presence of E-box variant only in the promoter region of CYCP3;1. **(H)** Bar chart of ChIP-qPCR showing enrichment levels of *CYCP3;1* and *CYCP3;2* and *CYCD1;1* in anti-GFP antibody pull-down immunoprecipitated (IP) samples compared to control (input) of *spt- 12*/*SPT::SPT:sYFP* inflorescences. The expression levels were normalised against *ACTIN7.* Error bars represent SD; *p<0.05; ns. signifies not statistically significant (unpaired Student’s *t*-test). n=3 biological replicates.

Additionally, SPT is involved in a complex interplay that coordinate the morphological auxin signals with the proliferative signal mediated by the hormone cytokinin (CK)(*13–15*) which plays antagonistic roles to auxin throughout plant development, including gynoecium patterning(*16, 17*). Accordingly, *spt* mutant styles are hypersensitive to CK applications, *i.e*., an increase of the frequency and severity of split styles is observed following 6-Benzylaminopurine (BAP) treatments(*18, 19*).

Despite the apical fusion of the gynoecium plays pivotal role in ensuring efficient fertilization and seed production, the molecular and cellular mechanisms that guide carpel fusion at the apical style remain elusive.

Coordination of proliferation and cell-division orientation has been proposed to facilitate division plane determination in proliferating tissues, in the root meristem(*20, 21*)

This scenario is consistent with a requirement for a tight regulation of cell-division activities during the apical fusion of the carpels, where we hypothesised a robust placement of cell division might underpin the final fusion of the apical-medial marginal tissue(*9*), where an auxin maximum picks and cytokinin signalling response is inhibited.

Combining genetic, molecular, and pharmacological experiments, we show that SPT directly and cell-autonomously represses the expression of *CYCLIN-P3;1* (*CYCP3;1*) and *CYCP3;2*(*22, 23*), specifically at the style, in an opposite fashion to the cell-proliferation signal provided by CKs, which induces their expression. We showed that CYCP3s activity is causative of the *spt* mutant bilateral style phenotype. Overexpressing CYCP3s lines displayed a significant number of unfused styles, resembling the *spt* split-style phenotype. Moreover, radial styles were partially restored in the *spt cycp3;1 cycp3;2* triple loss-of-function mutant, and hypersensitivity to CK was reduced, meaning CYCP3s play roles in controlling cell-proliferation via the CK signalling pathway. Furthermore, we showed that combining defects in cell-division orientation (by using loss-of-function mutants in key players involved with PPB function/assembly) with perturbation in proliferation (by a tissue-specific expression of a CK biosynthetic gene and via external CK applications), drastically impacts the apical fusion of the two carpels, *i.e.,* break of style radial symmetry. Thus, SPT regulates morphogenesis of the style by working cell-autonomously at the medial-apical cells, by simultaneously coordinating an auxin-centric incoherent feed-forward loop(*9*) with a CK-centric coherent feed-forward regulatory loop, involving CYCP3s, to add robustness to style morphogenesis.

## RESULTS

### *In vivo* regulation of gene expression by SPATULA

To reveal the genes and processes regulated by SPT in the apical style region of the *Arabidopsis* gynoecium, we performed chromatin immunoprecipitation (ChIP) followed by deep sequencing (Seq) using inflorescences material from the SPT complementation line (*spt-12/SPT*::*SPT*– *sYFP*)(*12*).

We selected a list of 6,731 high confidence candidate genes that consistently enriched in all three biological replicates using 0.001 FDR (false discovery rate).

At the top of the list, we found that SPT binds to its own promoter (Fig. 1d and Suppl. Table 1), which explains the high levels of SPT transcripts previously observed in an SPT overexpressing line(*24*) and in our *spt-12/SPT*::*SPT*–*sYFP* complementation line(*12*).

As further positive controls, we examined genes previously reported to interact with SPT genetically or functionally during gynoecium development, such as the bHLH TFs INDEHISCENT (IND)(*25*) and HECATEs (HECs)(*14, 26*), as well as CUP-COTYLEDON 2 (CUC2)(*27*) (Suppl. Fig. 1a-d and Suppl. Table 1). These genes were found to be bound by SPT in all three biological replicates, supporting the established model that these transcription factors function coordinatively(*28*).

To understand the biological processes controlled downstream of SPT, we analysed the top twenty Gene Ontology (GO) categories that were statistically enriched by ranking all 6,731 presumptive SPT direct target genes. We found the processes related to reproduction, development and multicellular organismal processes were significantly enriched and included a high number of genes, such as SPT, IND, HEC1,2, and CUC2 (Fig. 1e and Suppl. Table 1).

Other enriched GO terms highlighted roles for SPT in phosphorylation and phosphate-related metabolic processes, signal transduction, transport, catabolism and protein modification processes (Fig. 1e). These findings are consistent with previously identified roles of SPT in sugar-based post-translational modifications(*12*), as well as auxin and cytokinin signalling and transport(*13, 14, 29*).

The role of SPT in style radialization has recently been linked to the regulation of cell-division rate and orientation, specifically through the regulation of members of the D-type cyclin family (CYCDs), namely CYCD1;1 and CYCD3;3(*9*). Accordingly, we found enrichment of CYCD3;3 in two of the three biological replicates analysed (Suppl. Fig, 1e). However, neither loss-of-function nor overexpressing mutants of CYCDs have been reported to affect style morphogenesis on their own, suggesting that SPT regulates multiple aspects of cell-division control to ensure correct style morphogenesis.

To understand the underlying mechanisms involved in style morphology, we focused on presumptive downstream targets of SPT involved in cell-cycle and cell-division regulation (Suppl. Table 1). A previous study showed that among the cell-cycle-related genes regulated by IND, while CYCD1;1 was upregulated, a member of the P-type cyclin family (CYCPs), CYCP3;1, was found to be downregulated by IND(*7*). Interestingly, among the cell-cycle genes downstream of SPT (Suppl. Table 1) *CYCP3;2* was found to be a high confidence candidate target of SPT in all 3 biological replicates with a nearest peak summit at 1.58 kb upstream of transcription start site, and within 200bp vicinity of core hexanucleotide sequence i.e., G-box (CACGTG) previously reported to be bound by SPT(*30*) (Fig. 1f).

In Arabidopsis, CYCP3;2 and its closely related sister protein CYCP3;1 (hereafter together refereed as CYCP3s), have been recently proposed as positive regulators of cell proliferation(*22, 31*), although their roles in development and cell-cycle control remain largely unknown.

According with the presence of G-box elements in the promoter of both *CYCP3s* (Fig. 1g), we confirmed that both *CYCP3s* are direct target of SPT by performing q-PCR experiments from three independent biological ChIP material of *spt-12/SPT*::*SPT*–*sYFP* inflorescences (Fig. 1h). Our experiments also showed that another cyclin genetically linked to SPT and IND, CYCD1;1(*9*), is not a direct target of SPT (Fig. 1h).

Altogether our experiments shed light on the cellular and molecular roles of SPT activity and confirm that both *CYCP3s* are direct targets of SPT.

### SPATULA represses *CYCP3s* expression in a coherent feed-forward loop with CKs

To investigate the causative molecular mechanisms linking cell-division defects in the *spt* mutant bilateral style, we further characterised the roles of CYCP3s during style development. SPT has been reported to work as growth repressor in roots and cotyledons(*32–34*). Accordingly, in the style, SPT represses the CK output(*14*), which has a proliferative effect on the abaxial marginal gynoecium tissues(*19*) (Fig. 1a). To pattern the style, SPT finetunes the auxin/CK crosstalk which have opposite effects on style development(*15*): while blocking auxin transport by NPA applications lead to a thinner and longer wild-type style without inducing ectopic growth, CK applications results in wider and shorter style and promotes ectopic proliferation (Fig. 1a-c). Accordingly, *spt* bilateral style frequency and complexity is highly increased by CK applications(*14*), strongly suggesting that SPT may control style morphology by repressing proliferation in an opposite manner to cytokinin.

Because CYCP3s were also found among the downstream targets of the CK response regulators in the shoot apical meristem(*35*), we hypothesised that CYCP3s could be transcriptionally regulated by SPT and CK in an antagonistic way, *i.e.*, downregulated by SPT and upregulated by CK at the style region.

We constructed and analysed *CYCP3s* transcriptional GUS-fusion (*pCYCP3;1:GUS* and *pCYCP3;2:GUS*) in WT and *spt-12* backgrounds. In line with the hypothesised repressive activity mediated by SPT, neither *CYCP3s* were expressed in the wild-type style (Fig. 2a,b). Interestingly, *CYCP3;1* was expressed in the adaxial tissue of the bilateral ovary, the endocarp-*a*, from stage 9 of gynoecium development onwards (Fig. 2a and Suppl. Fig. 2a). *CYCP3;2* was not expressed in any tissue of the developing gynoecium (Fig. 2b) until very later in development, after the patterning of the style is completed (Suppl. Fig. 2b). Optical microscopy analysis revealed that both *CYCP3s* were upregulated in the *spt* mutant background, specifically in the medio-apical region of the style, where the unfused carpels break radial symmetry (Fig. 2a,b).

**Fig. 2.**
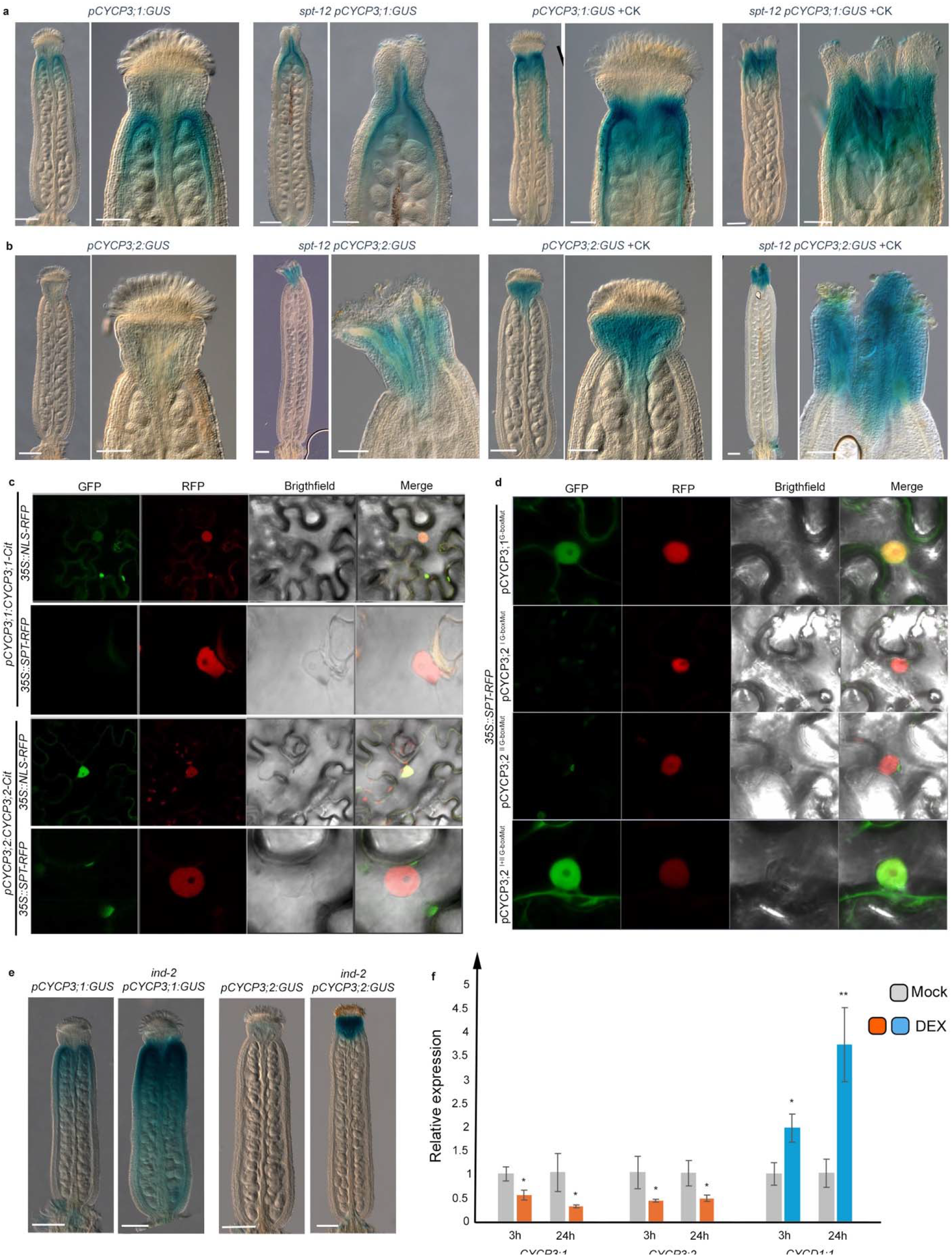
SPATULA and Cytokinin regulate CYCP3s expression in an antagonistic fashion. (A, **B)** Light microscope images of GUS-stained gynoecia of p*CYCP3;1:GUS* (a) and p*CYCP3;2:GUS* (b) in wildtype Col-0 and *spt-12* mutant backgrounds with no treatment (a and b, left-hand side) and with CK (50 μM BAP) treatment (a and b, the right-hand side). Scale bars represent 200 μm (full-size gynoecia) and 100 μm (style magnifications). n=3 biological replicates and ∼25-30 gynoecia were analysed for each genotype and treatment. **(C)** Confocal images of tobacco leaf cells co-infiltrated with either *pCYCP3;1:CYCP3;1-Cit* or *pCYCP3;2:CYCP3;2-Cit* plus *35S::SPT-RFP* or *35S::NLS-RFP* as depicted on the panels. GFP, RFP, Brightfield and merged images are shown. n=3 biological replicates. Note, *CYCP3s* are expressed only in the absence of SPT. **(D)** Confocal images of tobacco leaf cells co-infiltrated with *pCYCP3;1^G-mut^:CYCP3;1:Cit* and *35S::SPT-RFP* (top panels); *35S::SPT-RFP* co-expressed with either single mutated versions of *pCYCP3;2* (*pCYCP3;2^I^ ^G-boxMut^:CYCP3;2-Cit and pCYCP3;2 ^II^ ^G-boxMut^:CYCP3;2-Cit)* (middle panels) or double mutated version (*pCYCP3;2^I+II^ ^G-boxMut^:CYCP3;2-Cit)* (bottom panels). GFP, RFP, Brightfield and merged images are shown. n=3 biological replicates. **(E)** Light microscope images of GUS-stained gynoecia of p*CYCP3;1:GUS* and p*CYCP3;2:GUS* in wildtype Col-0 and *ind-2* mutant backgrounds. Scale bars represent 200 μm. n=3 biological replicates and ∼25-30 gynoecia were analysed for each genotype. **(F)** Bar chart of qRT-PCR of *CYCP3;1*, *CYCP3;2* (orange bars) and *CYCD1;1* (blue bars) expression levels after 3h and 24h of DEX induction (orange and blue bars) of *35S::IND:GR* line compared to Mock (grey bars), normalised against *UBIQUITIN10*. n=3 biological replicates with 10 seedlings per replicate. Error bars represent SD; *p<0.05, **p<0.001 (unpaired Student’s *t*-test).

In addition, we performed co-expression experiments in tobacco leaves for both CYCP3s genomic sequence driven by their native promoters and fused to CITRINE (Cit) (*pCYCP3;1:CYCP3;1-Cit* and *pCYCP3;2:CYCP3;2-Cit*), co-expressed with either *35S::SPT-RFP* or a *35::NLS-RFP* (Fig. 2c). We observed that CYCP3s were expressed in tobacco epidermal cells and localised in both nucleus and cytoplasm, only when co-expressed with *35::NLS-RFP* construct. By contrast, the co-expression of *35S::SPT-RFP* with either *pCYCP3s:CYCP3s:Cit* caused the elimination of CYCP3s expression (Fig. 2c). To prove the downregulation mediated by SPT was operating at the transcriptional level, we mutated the single and double G-box elements included in *CYCP3;1* and *CYCP3;2* promoters, respectively (Fig. 1g, Fig. 2d). Mutation of the single G-box element in *CYCP3;1* promoter (*pCYCP3;1^G-mut^:CYCP3;1:Cit*) co-expressed with *35S::SPT-RFP* rescued the expression of CYCP3;1 (Fig. 2d). For CYCP3;2, mutation of each single G-box elements (*pCYCP3;2^I^ ^G-boxMut^:CYCP3;2:Cit and pCYCP3;2^II^ ^G-boxMut^:CYCP3;2:Cit*) was not sufficient to do so, while the simultaneous mutation in both G-boxes (*pCYCP3;2^I+II^ ^G-boxMut^:CYCP3;2:Cit)* enabled the recovery of CYCP3;2 when co-expressed with *35S::SPT-RFP* in tobacco leaves (Fig. 2d).

Furthermore, SPT interacts genetically and physically with another bHLH transcription factor INDEHISCENT (IND), which binds to the E-Box variant (CACGCG)(*36*) (Fig. 1g). According to the synergistic activity of SPT and IND, *CYCP3s* expression was found to be upregulated in *ind-2* apical style *in vivo* (Fig. 2e), and down-regulated *in vitro* in seedlings of the *35S::IND:GR* overexpression line followed by 3h and 24h of IND induction by Dexamethasone (DEX) treatments (Fig. 2f).

Altogether these data demonstrate that SPT and IND both downregulate *CYCP3;1* and *CYCP3;2* expression.

In line with several CK-binding sites (GARP AGATT(T/C)(*37*) present in the promoters of *CYCP3s* (Fig. 1g), next we tested the role of CKs on the *in vivo* expression of *CYCP3s* at the style region. CK treatments followed by GUS staining of *pCYCP3;1:GUS* and *pCYCP3;2:GUS* reporters revealed that both *CYCP3s* were ectopically upregulated, specifically at the gynoecium apex, following exogenous hormonal applications (Fig. 2a,b), mimicking the expression seen in the *ind* and *spt* single backgrounds without CK treatments. Furthermore, *CYCP3s* ectopic upregulation in the style region was further enhanced in the *spt* background *in vivo* by CK treatments, supporting an antagonistic and independent effect of SPT and CK on the control of *CYCP3s* expression(*14, 19*) (Fig. 2a,b).

To further test the CK-mediated control of CYCP3s expression, we used an inducible constitutive active form of a B-type Arabidopsis Response Regulators (ARR-B), ARR1, (*35S::ARR1*Δ*DDDK:GR*)(*38, 39*), a positive signalling component of the CK pathway, and compared DEX-treated inflorescences of *35S::ARR1*Δ*DDDK:GR pCYCP3;2:GUS* and *35S::ARR1*Δ*DDDK:GR pCYCP3;1:GUS* to mock-treated controls (Suppl. Fig. 2c,d). These experiments revealed that *CYCP3;2* expression was ectopically induced in the endocarp-*a* (mimicking the expression of *CYCP3;1*) and in the style by the CK signalling, while CYCP3;1 was not affected by the constitutive activation of ARR1. This *in vivo* analysis showed that induction of ARR1 is sufficient to upregulate at least *CYCP3;2* expression in the gynoecium tissues (Suppl. Fig. 2d), while CYCP3;1 might be regulated by other type-B ARRs.

Altogether, our data provide the first direct *in vivo* evidence for an opposite transcriptional regulation of two potential core players of the cell-cycle by SPT and CK during organ development, in a coherent forward loop type-II(*40*).

### Ectopic activity of CYCP3;1 and CYCP3;2 break radial symmetry at the gynoecium apex

To understand whether CYCP3s function mediate cell proliferation triggered by CK, we tested whether a double *cycp3s* loss-of-function mutant display resistance to CK applications. We constructed a double CRISPR-Cas9 mutants for both CYCP3s (Suppl. Fig.3a-d) and treated it with exogenous CK applications. In this double *cycp3s* CRISPR mutant, guides were directed upstream of the cyclin-box motif (necessary for Cyclins binding to CDKs) which is contained in the first exon of both CYCP3s (Suppl. Fig.3a,b). The premature stop codon presumably produces truncated protein forms, which we predict to change CYCP3;1 from 220 amino acid (aa) residues into an allele we named *cycp3;1-1* of 96 aa, and CYCP3;2 from 230 aa to the allele *cycp3;2-1* of 90 aa (Suppl. Fig.3c,d). SEM analysis of wild-type and *cycp3;1 cycp3;2* double mutant gynoecia treated with mock and CK showed absence of ectopic growth from the medial tissues of the *cycp3s* CRISPR double mutant ovary and style (Fig. 3a-c), suggesting that CK-induced proliferation output in the gynoecium requires CYCP3s function.

**Fig. 3.**
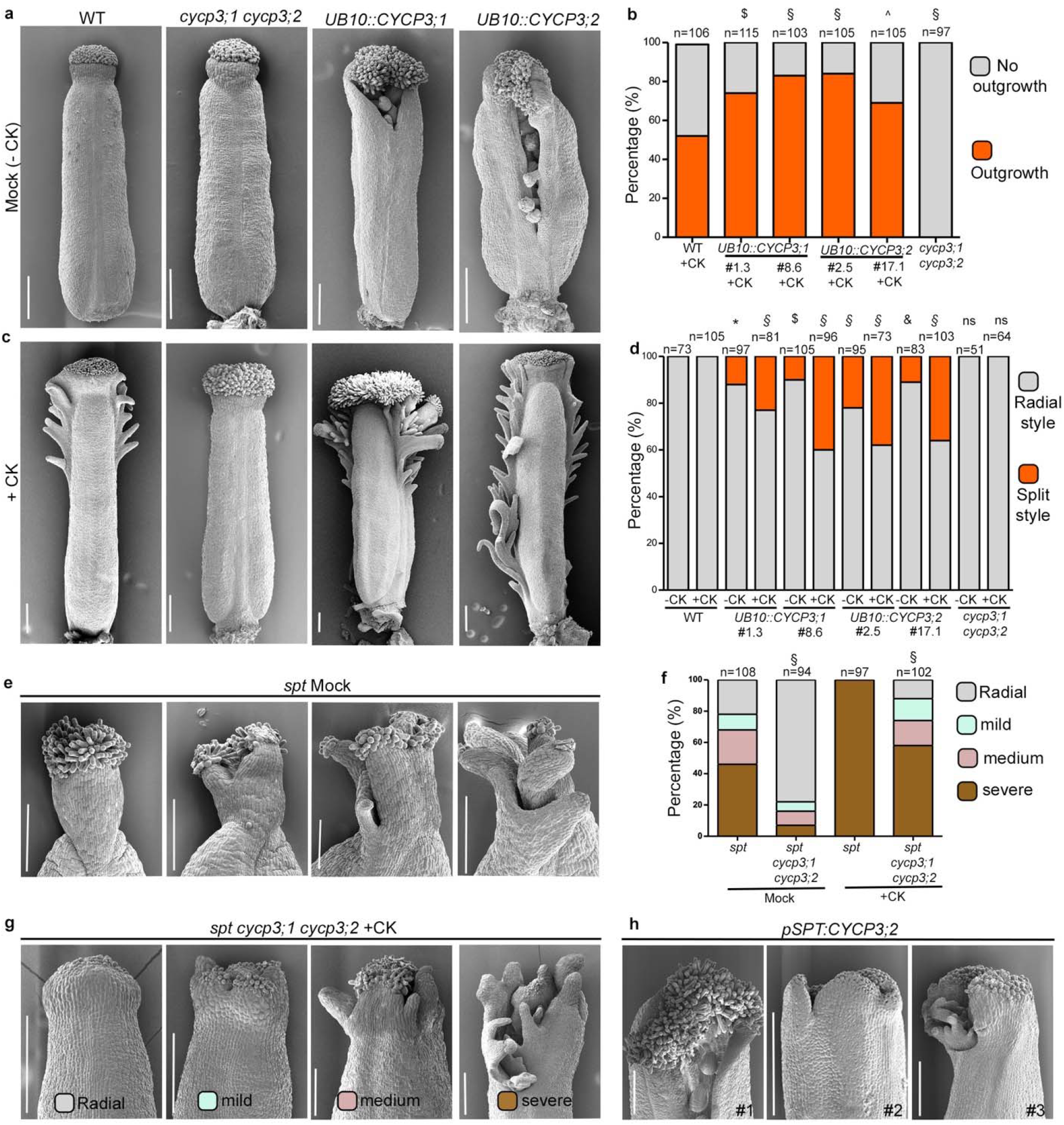
CYCP3s function is necessary and sufficient to control style radial symmetry. (A, C) SEM images of mock (-CK) (a) and CK (+CK) (c) treated gynoecia of wildtype Col-0, double *cycp3s* CRISPR mutant (*cycp3;1cycp3;2*), and a representative homozygous overexpressing lines of *UB10::CYCP3;1:Myc* (#1.3) and *UB10::CYCP3;2:Myc* (#2.5). Scale bars are 200 μm. **(B)** Bar chart of quantification (percentage) of ectopic outgrowths phenotype in +CK treated gynoecia of wildtype Col-0, *cycp3;1cycp3;2*, and two independent overexpressing homozygous lines of *UB10::CYCP3;1:Myc* (#1.3 and #8.6) and *UB10::CYCP3;2:Myc* (#2.5 and #17.1). Grey bars represent ‘no outgrowths’ and orange bars represent ‘outgrowths’. 2x2 contingency table followed by Fisher’s exact Chi^2 test was used to compare phenotypic classes. Two tailed *p* values are as follows: Col-0 CK vs #1.3 CK, p=0.0032 ($), Col-0 CK vs #8.6, Col-0 CK vs #2.5 and Col-0 CK vs *cycp3;1 cycp3;2*, p<0.00001 (§), and Col-0 CK vs #17.1, p=0.04 (^). Three biological replicates have been performed; results of one representative biological replicate are plotted. Number of samples (n) analysed for each genotype/treatment are written on top of bars. **(D)** Bar chart of quantification (percentage) of radial (grey bars) and split (orange bars) phenotypes of -CK and +CK treated gynoecia of wildtype Col-0, *cycp3;1cycp3;2*, and two independent overexpressing homozygous lines of *UB10::CYCP3;1:Myc* (#1.3 and #8.6) and *UB10::CYCP3;2:Myc* (#2.5 and #17.1). 2x2 contingency table followed by Fisher’s exact Chi^2 test was used to compare phenotypic classes. Two tailed *p* values are as follow: Col-0 mock vs #1.3 mock, p=0.0003 (*); Col-0 mock vs #8.6 mock, p<0.006 ($); Col-0 mock vs #2.5 mock, p<0.00001 (§); Col-0 mock vs #17.1, p<0.0007 (&) Col-0 mock vs *cycp3;1 cycp3;2* mock p=1 (ns=non-significant); Col-0 Ck vs #1.3 CK, Col-0 Ck vs #8.6 CK, Col-0 Ck vs #2.5 CK, Col-0 CK vs #17.1 CK, p<0.00001 (§) and Col-0 CK vs *cycp3;1 cycp3;2* CK, p=1 (ns=non-significant). Three biological replicates have been performed; results of one representative biological replicate are plotted. Number of samples (n) analysed for each genotype/treatment is written on top of bars. **(E)** SEM images of styles of *spt* mock (internal control) gynoecia showing 4 different categories of phenotypes, including radial (grey), mild split (blue), medium split (pink) and severe split (brown). Scale bars are 200µm. **(F)** Bar chart showing percentage of radial and split (mild, medium, severe) phenotypes of mock and CK treated (+CK) *spt* (internal control) and *spt cycp3;1 cycp3;2* gynoecia. 2x2 contingency table followed by Fisher’s exact Chi^2 test was used to compare phenotypic classes. Two tailed *p* values are as follow: *spt* mock vs *spt cycp3;1 cycp3;2* mock, and *spt* +CK vs *spt cycp3;1 cycp3;2* +CK, p<0.00001 (§). Three biological replicates have been performed; results of one representative biological replicate are plotted. Number of samples (n) analysed for each genotype/treatment is written on top of bars. **(G)** SEM images of styles of *spt cycp3;1 cycp3;2* gynoecia treated with CK (+CK) showing 4 different categories of phenotypes, including radial (grey), mild split (blue), medium split (pink) and severe split (brown). Scale bars are 200µm. **(H)** SEM images of styles of gynoecia of 3 independent T1 lines of *pSPT:CYCP3;2:HA* (#1, #2, #3). Scale bars are 200µm.

To test whether CYCP3s function is sufficient to break radial symmetry at the style, we overexpressed CYCP3;1 and CYCP3;2 (*UB10::CYCP3;1:Myc* and *UB10::CYCP3;2:Myc*) (Suppl. Fig. 3e,f) and analysed their gynoecia by SEM. A population of *CYCP3;1* and *CYCP3;2* overexpressing lines showed a consistent split style phenotype (Fig. 3a,d). In line with a positive role in CK signalling, CK applications increased the frequency of the split-style observed in the *UB10::CYCP3s:Myc* lines to 44%, while it had no effect in the wild-type background (Fig. 3a,d and Fig. 4c) as well as the ectopic growth arising from the ovary (Fig. 3b,c). The hypersensitivity of CYCP3s to CK corroborate a functional role for CYCP3s in controlling cell proliferation and response to CK at the style region and control of style morphology.

**Fig. 4.**
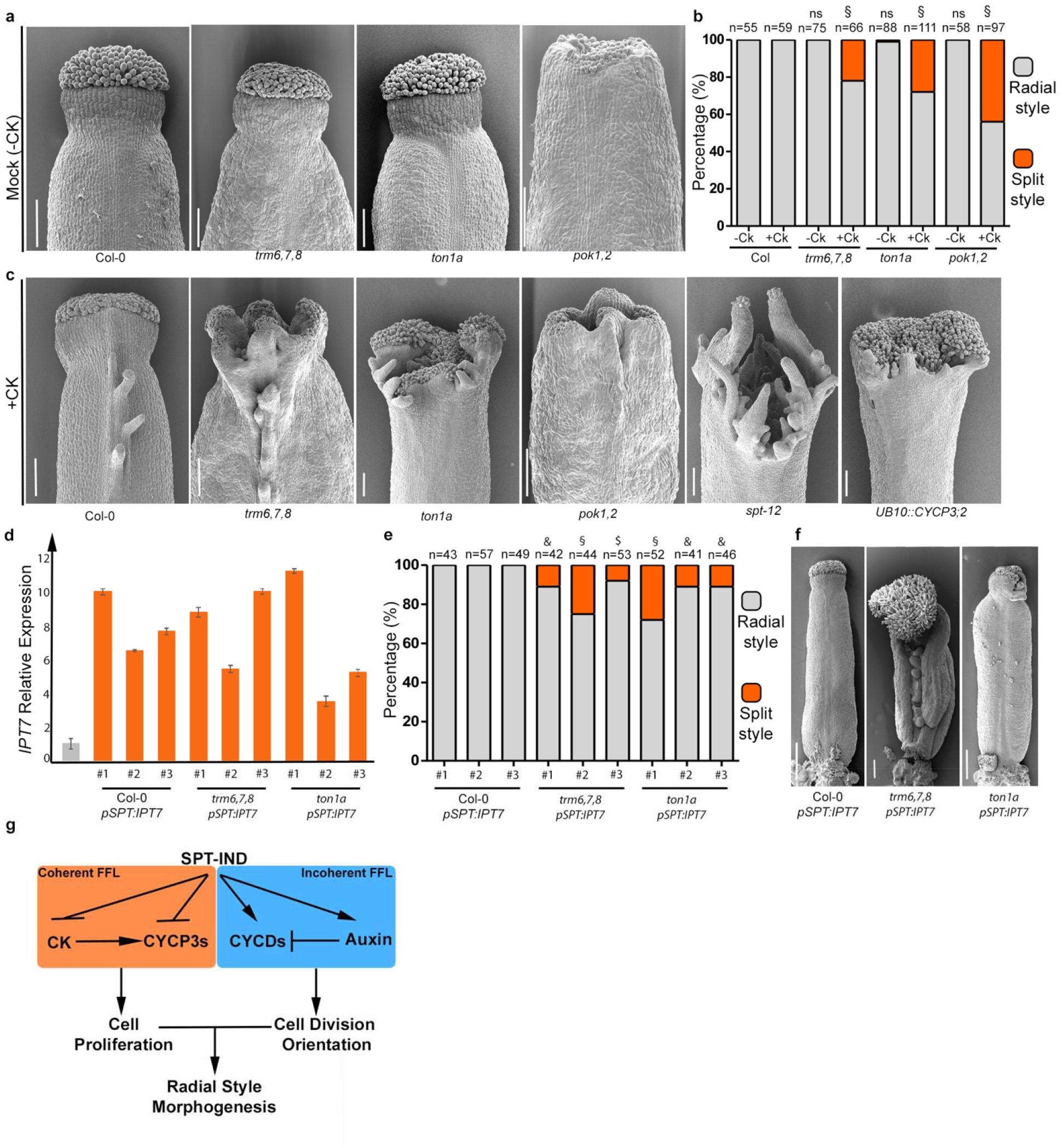
Cytokinin applications impact style morphology in cell-division orientation mutants. (A) SEM images of mock (-CK) treated gynoecia of wildtype (Col-0) and mutants in cell-division orientation (*trm6,7,8, ton1a, pok1,2*) showing radial styles. Scale bars represent 100µm. **(B)** Bar chart showing quantification (percentage) of radial (grey bars) and split (orange bars) style phenotypes of -CK and +CK treated gynoecia of Col-0, *trm6,7,8, ton1a,* and *pok1,2*. 2x2 contingency table followed by Fisher’s exact Chi^2 test was used to compare phenotypic classes. Two tailed *p* values are as follow: Col-0 mock vs *trm6,7,8* mock, Col-0 mock vs *ton1a* mock, and Col-0 mock vs *pok1,2* mock, *p* = 1 (ns=non-significant); Col-0 CK vs *trm6,7,8* CK, Col-0 CK vs *ton1a* CK, Col-0 CK vs *pok1,2* CK, p<0.0001 (§). Three biological replicates have been performed; results of one representative biological replicate are plotted. Number of samples (n) analysed for each genotype/treatment is written on top of bars. (**C)** SEM images of CK treated (+CK) Col- 0, *trm6,7,8, ton1a, pok1,2, spt-12* and *UB10::CYCP3;2* gynoecia. Scale bars are 100µm**. (D)** Bar chart of qRT-PCR showing relative expression levels of CK biosynthesis gene *IPT7* in 3 independent T1 lines of *pSPT:IPT7* in Col-0, *trm6,7,8*, and *ton1a* backgrounds. Expression levels were normalised against *UBIQUITIN10*. The experiment was performed once on three independent transgenic lines per construct, with four technical repeats. Values shown are means ± SEM. **(E)** Bar chart showing quantification (percentage) of radial (grey bars) and split (orange bars) style phenotypes of Col-0, *trm6,7,8* and *ton1a* gynoecia transformed with *pSPT:IPT7* construct. Three independent T1 lines for each background are plotted. Number of gynoecia analysed for each genotype are shown on top of respective bars. 2x2 contingency table followed by Fisher’s exact Chi^2 test was used to compare phenotypic classes. Two tailed *p* values are as follow: Col-0 *pSPT:IPT7* vs *trm6,7,8 pSPT:IPT7* line *#1*, and Col-0 *pSPT:IPT7 vs ton1a* lines #2 and #3, *p*<0.0007(&); Col-0 *pSPT:IPT7* vs *trm6,7,8 pSPT:IPT7* line *#2,* and Col-0 *pSPT:IPT7* vs *ton1a pSPT:IPT7* line *#1, p*<0.00001 (§), Col-0 *pSPT:IPT7* vs *trm6,7,8 pSPT:IPT7* line #3, p<0.006 ($). **(F)** SEM images of representative gynoecia from one independent transgenic T1 line of Col-0, *ton1a,* and *trm6,7,8* transformed with *pSPT:IPT7* construct. Scale bars are 200µm. **(G)** Schematic model showing the regulatory network of *CYCP3s* and *CYCDs* expression by bHLH TFs (SPT-IND) and hormones CK and auxin forming a coherent and incoherent feed forward loop (FFP), respectively, to regulate radial style morphogenesis.

Next, we asked whether the ectopic *CYCP3s* expression observed in the *spt* split-style was causative of its bilateral phenotype and hypersensitivity of the *spt* style phenotype to CK applications. We analysed *spt cycp3;1 cycp3;2* triple mutant gynoecia by SEM and quantified the presence of bilateral vs radial style compared to a segregating *spt* control. The analysis showed that proper fusion of the style displaying radial symmetry was largely restored in the *spt cycp3;1 cycp3;2* mutant gynoecia (78% radial and 22% bilateral styles) compared to the *spt* segregating control (22% radial and 78% bilateral styles) (Fig. 3e-f). Furthermore, CK application experiments showed that the gynoecia apices of the triple *spt cycp3;1 cycp3;2* were partially resistant to hormonal treatments compared to the *spt* single mutant. In these experiments we divided the split-style phenotype in three further categories to account for the severity and complexity of the cleft, as severe, medium, and mild (Fig. 3e-g). As previously shown, *spt* style phenotype increased drastically by CK applications, leading to 100% of samples showing a severe split style compared to mock-treated *spt* mutants (Fig. 3f and Fig. 4c).

Mock-treated *spt cycp3;1 cycp3;2* gynoecia showed a vast increase in radial, fused style (78%) and a reduced frequency of severe (7%), medium (9%), and mild (6%) split-style phenotypes (Fig. 3f). Although CK treatments still raised the frequency of severe (58%), medium (16%), and mild (14%) split styles, 12% of radial styles were still present after the treatment of *spt cycp3;1 cycp3;2* gynoecia, which is never the case for the *spt* single mutant (Fig. 3f,g). This data demonstrates that CYCP3s promote ectopic cell-proliferation in the *spt* mutant background via promoting the CK mediated cell-division input and style morphology.

Lastly, to test whether ectopic CYCP3s function could break style radial symmetry by working cell-autonomously in the SPT-expression domain, we analysed *pSPT:CYCP3;2:HA* transgenic lines that showed overexpression of CYCP3;2 transcript (Suppl. Fig. 3g) and observed split-style phenotype that resembles that of both *spt* and *UB10::CYCP3s* mutants (Fig. 3h). This corroborates a model where SPT represses *CYCP3;2* (and *CYCP3;1*) expression cell-autonomously to fuse the apical carpels.

Altogether, our data show that the activity of CYCP3s is sufficient to break radial symmetry at the gynoecium apex and necessary to confer sensitivity to CK when SPT function is missing.

### Increasing CK levels affect symmetry establishment in mutants impaired in cell-division orientation

SPT promotes accumulation of auxin maxima foci, which in turn are important to maintain division orientation in the periclinal direction(*9*) to form a radial style(*2, 3*). Division plane orientation in plants is established before mitosis and must be maintained throughout mitosis and cytokinesis(*41*). This process is often thought to occur when the preprophase band (PPB) forms in G2 phase of the cell-cycle(*41–43*) although the current scenario assigns the PPB a marginal role in determining cell-division orientation, compared to the past view: the PPB would add robustness to the selection of the right division angle, rather than being a key determinant of cell-division orientation *per se*(*43*).

Key components of the PPB establishment machinery such as TONNEAU 1A (TON1A)(*44*), the TON1 Recruiting Motif 6,7,8 (TRM6,7,8)(*43*) proteins, and PPB maintenance PHRAGMOPLAST ORIENTING KINESIN 1 (POK1) and POK2(*45*) were found among the presumptive SPT downstream targets (Table 1). Notably, although the PPB still form in the *spt* background, its orientation is miss-placed at the mutant developing style(*9*).

How cell-division orientation governs carpels apical fusion and style shape is unknown. To understand whether correct orientation of cell-division is required for style radialization, we tested whether mutants impaired in cell-division orientation would display a split-style phenotype, similar to *spt*. To test the importance of cell-division orientation in style development, we analysed by scanning electron microscopy (SEM) gynoecia of mutants in key components of the microtubule-dependent mitotic structures important for the cell-cycle: the PPB establishment, guided by TON1a and TRMs 6,7,8, (*ton1a-1* and *trm6, trm7 and trm8*)(*43, 44*), and PPB maintenance guided by POK1 and POK2 (*pok1,2*)(*45*). None of the aforementioned mutants showed significant defects in style development (Fig. 4a,b). This suggests that either cell-division orientation is not essential *per se* for radial style morphogenesis, or a synergistic layer of control adds robustness to keep the orientation of newly forming cell walls perpendicular to the direction of growth(*9*), which ultimately guarantees apical-basal anisotropic growth(*10*).

Because SPT promotes auxin accumulation and cell-division orientation(*2, 3, 9*) and represses CK signalling(*14*), dampening the mitotic potential at the marginal tissue(*19*), we asked whether augmenting CK levels by exogenous applications in cell-division orientation mutants would break radial symmetry and mimic the *spt* split style. SEM analysis of *ton1a*, *trm6,7,8* gynoecia of inflorescences treated with CK and mock, showed that CK applications uncovered a never-seen before split-phenotype of these mutants, leading to a significant percentage of split styles recovered in *ton1a*, *trm6,7,8* and *pok1,2* (Fig. 4a-c). Moreover, these severe split-style phenotypes displayed augmented complexity of their distal gynoecia apices, similar to *spt* gynoecia treated with CK (Fig. 4c). In addition, tissue-specific expression of the CK biosynthetic gene *IPT7*(*46*) driven by the SPT promoter (*pSPT:IPT7*) (Fig. 4d) showed increased frequency of split-style phenotypes observed in *ton1a* and *trm6,7,8* mutant backgrounds but not in the wild-type control (Fig.4 e,f), yielding similar results to those obtained with CK exogenous applications (Fig. 4a-c). Altogether our results demonstrate a coordination between cell proliferation and division orientation to fuse the apical style.

In conclusion, our data support a model where a key regulator of style development, SPT, provides robustness to the placement of the cell-division angle by a local fine-tuning of cell-cycle progression and proliferation mediated via the control of two families of cyclins: CYCD1;1 and CYCD3;3 via auxin(*9*) and CYCP3;1 and CYCP3,2 via CK (Fig. 4g), ultimately important for shaping radial symmetry at the style region.

## DISCUSSION

Altogether, our results demonstrate a role for the presumptive cell-cycle regulators CYCP3;1 and CYCP3;2 in controlling radial symmetry establishment at the gynoecium apex, a critical step required for the final fusion of the two carpels in *Arabidopsis* reproductive development.

SPT has long been implicated in the antagonistic crosstalk between auxin and cytokinin (CK), but the downstream activities of this network remain unclear.

We demonstrated that the antagonistic roles of the bHLH transcription factor SPT and CK control the expression of *CYCP3s* in opposite ways. SPT directly represses *CYCP3*s expression in the style region, while CK promotes their expression, forming a coherent feed-forward loop (FFL) type II(*40*). This opposing regulation of *CYCP3s* by SPT and CK aligns with previous findings that CYCP3s are positive targets of CK signalling in the shoot apical meristem(*47*) and negatively regulated by the SPT-interacting factor, IND(*7*).

Coherent FFLs have been suggested to add robustness to signalling networks by making downstream nodes more resistant to perturbations compared to single incoherent FFLs(*48*) . We previously showed that SPT (and IND) also participate in an incoherent FFL with auxin to regulate the expression of D-type cyclins, potentially coordinating G1 cell-cycle progression and the orientation of cell division(*9*). Interestingly, while incoherent FFLs accelerate the response of target genes, coherent FFLs tend to delay them(*49*). This suggests a model where SPT fine-tunes cell-division activities by interacting with auxin and CK signalling through these two types of FFLs (Fig. 4g).

Carpel fusion is essential for efficient fertilization and seed production. Growth analyses of the wild-type gynoecium reveal that strong anisotropic growth in the apical-basal direction drives the development of epidermal tissue in the style region(*10, 11*). Cell clones in the style expand longitudinally and divide in the transverse anticlinal direction(*10*). Our recent analysis of the *spt* mutant bilateral style revealed abnormalities in cell-division orientation control at the onset of the cleft, where an auxin maximum accumulates to drive carpel fusion(*9*). To test this model, we investigated whether mutants with aberrant cell-division orientation, such as *ton1a* and *trm6,7,8*, displayed defects in style morphology. Our genetic analysis showed that defects in cell-division orientation alone are insufficient to disrupt radial symmetry in the style, suggesting an additional layer of control that enhances robustness. We found that increasing CK levels in PPB-defective mutants, via exogenous CK application or endogenous CK production by the *IPT7* gene, resulted in severe style defects similar to the *spt* phenotype. This confirms that CK impacts radial style formation in cell-division mutant backgrounds by at least influencing cell proliferation. It also highlights a link between CK action and key players in PPB establishment, such as TON1a(*44*) and TRM6,7,8(*43*), which add robustness to cell-division plane placement during style radial patterning. This may reflect a specific role for CK in carpel fusion control, where proper cell-division orientation requires additional regulation to ensure successful carpel fusion and reproductive fitness.

We demonstrated that CYCP3s function promote CK-mediated cell proliferation, as CRISPR mutants with *cycp3;1-1 cycp3;2-1* loss-of-function and CYCP3-overexpressing lines were resistant and hypersensitive to CK applications, respectively, as measured by the extent of outgrowths in the ovary region (Fig. 3b). However, how CYCP3s control the cell cycle remains unclear. If CYCP3s act downstream of CK, their function likely occurs during the G2 phase of the cell cycle, where CK promotes cell proliferation in the shoot apical meristem. This would also be compatible with a direct or indirect role for CYCP3s in controlling PPB formation, which also occurs in the G2 phase(*41, 42*).

In addition to proliferation control, CYCP3s may affect other processes, such as controlling the orientation of cortical or cytoplasmic microtubules (MTs). Recent research has shown that a radial arrangement of cytoplasmic MTs precedes PPB formation and enables cells to sense their geometry, facilitating symmetric division and the precise partitioning of cell volume into two daughter cells(*50*).

Since increasing cell proliferation in cell-division orientation mutants is sufficient to disrupt radial symmetry in the style, this suggests that CK promotes growth and/or cell expansion in the medio- lateral direction, antagonizing the auxin-mediated apical-basal style expansion. This may shift growth from anisotropic to isotropic, breaking the standard rules of growth and division at the gynoecium apex(*10, 11*). Supporting this, CK has been shown to regulate directional cell expansion/growth via cortical microtubule rearrangement(*51*).

A dual role for CYCPs is consistent with the characterization of CYCP homologs in the unicellular species *Trypanosoma brucei* (CYC2, CYC4, CYC5, and CYC7). When these cyclins are silenced, cells arrest in the G1 phase, and incorrect microtubule assembly leads to posterior axis bifurcation(*52*).

Taken together with our previous findings^9^, this work shows that SPT integrates fundamental cellular processes—including cell-division orientation, G1 phase progression, and cell proliferation—to orchestrate the morphogenesis of a radial style (Fig. 4g).

## MATERIALS AND METHODS

### Plant materials and growth conditions

The following mutants in the wildtype ecotype Columbia (Col-0) were used in the study: *spt- 12*(*1*), *ind-2*(*53*), *trm6,7,8*(*43*), *ton1a*(*44*), *pok1,2*(*45*)*, 35S::ARR1*Δ*DDK:GR*(*39*) and *35S::IND:GR*(*54*). *spt-12*, *ind-2* and *35S::ARR1*Δ*DDK:GR* mutants were crossed with transcriptional fusion lines *pCYCP3;1:GUS* and *pCYCP3;2:GUS*. Homozygous *cycp3;2*-1 mutant was crossed to *spt-12* mutant and the resulting double mutant was crossed to homozygous *cycp3;1-1* mutant to obtain a *spt cycp3;1 cycp3;2* triple mutant as well as a segregating *cycp3;1 cycp3;2* double mutant. The SPT complementation line *spt-12/SPT*::*SPT*–*sYFP* line used for ChIP-Seq analysis in this study was described previously(*12*). Plants were grown in JOHN INNES F2 STARTER soil mix (100% Peat, 4Kg/M3 dolomitic limestone, 1.2Kg/M3 osmocote start) in controlled environment room (CER) set at 22°C, with 80% relative humidity, in long day conditions (16h light/8h dark).

### Gynoecium and seedling treatments

To test the phenotypical effect of Cytokinin (CK) on gynoecia, treatments of Col-0, *UB10::CYCP3;1:Myc, UB10::CYCP3;2:Myc, cycp3;1 cycp3;2, spt-12, spt cycp3;1 cycp3;2, trm6,7,8, ton1a* and *pok1;2* plants were performed by spraying with 50 μM 6-Benzylaminopurine (BA Merck) and mock (NaOH), twice a week (4 times in total). First spray was done one week after bolting, and samples were collected after 4 days of last spray. To test the phenotypical effect of Naphthylphthalamic acid (NPA) treatment on Col-0 style, plants were treated with 100 μM NPA (Sigma) and mock (Absolute ethanol, Sigma), twice a day (morning and evening); samples were collected after 1 week of treatment. For expression analysis via GUS staining, homozygous plants of *pCYCP3;1:GUS*, *pCYCP3;2:GUS, spt-12 pCYCP3;1:GUS, spt-12 pCYCP3;2:GUS, ind-2 pCYCP3;1:GUS,* and *ind-2 pCYCP3;2:GUS,* were sprayed with 50 μM BAP for two consecutive mornings and samples for RNA extractions were collected the 3^rd^ morning. For Dex treatments of *35S::ARR1*Δ*DDK::GR pCYCP3;1:GUS* (F1) *and 35S::ARR1*Δ*DDK::GR pCYCP3;2:GUS* (F1) for GUS staining analysis, plants were sprayed with 10μM dexamethasone (DEX) (Sigma Aldrich, #218928) for two consecutive mornings, followed by samples collection on 3^rd^ morning. Silwet L-77 (0.015%) was used in all spray treatments. For DEX treatment of *35S::IND:GR* line for qRT-PCR analysis, seedlings were grown vertically on Murashige and Skoog (MS) media for 5 days before shifting to DEX (10μM) or Mock (DMSO, Sigma) containing MS plates for 3hrs and 24hrs time periods. Samples for RNA extractions were collected after 3hrs and 24hrs of mock and DEX treatments.

### DNA Constructs

#### UB10-driven overexpressing CYCP3s lines

Constructs of *UB10::CYCP3;1:Myc* and *UB10::CYCP3;2:Myc* were assembled using Golden Gate modular cloning method(*55*) as follows: The genomic coding sequence of CYCP3;1 (777bp) and CYCP3;2 (816bp) were amplified from genomic Col-0 DNA using the primers pairs CYCP3;1_gORF_F/CYCP3;1_gORF_R and CYCP3;1_gORF_F/CYCP3;1_gORF_R, respectively (no stop codon was included in the reverse primers). A proofreading Taq (Q5, NEB) was used for the PCR reaction. The amplification products were run on a 0.8% agarose gel to assess their size. The rest of the PCR reaction was purified using the (QIAGEN) and used in combination with the L0 Golden Gate vector pICSL01005 using Bbs1 (Sigma-Aldrich) and T4 ligase (Sigma-Aldrich). Constructs were transformed to *Escherichia coli* (DH5α) competent cells and positive colonies were selected on Spectinomycin (Spec^R^) (Sigma-Aldrich) LB plates. Plasmid DNA was extracted using NucleoSpin^®^ Plasmid kit by MACHEREY-NAGEL, enzymatic digestions were done using BsaI (NEB) and absence of mutations was confirmed by Eurofins sequencing (OVERNIGHT Mix2Seq kit). To generate level 1 constructs: amplicons of Level 0 modules, including the *UB10* promoter (pICSL12015), L0 vectors of pICSL01005_*CYCP3;1* and pICSL01005_*CYCP3;2*, C-terminal *Myc* tag (pICSL50010) and *NOS* terminator (pICH41421) were combinatorically assembled into Level 1 acceptor backbone (pICH47742) using a digestion and ligation (dig-lig) protocol with the type II restriction enzyme *Bsa*I and T4 DNA ligase. Both constructs were transformed to *Escherichia coli* (DH5α) strain. The transformed cells were selected on LB medium (Carb^R^), and incubated overnight at 37 °C. Cultured colonies were screened by mini-prep followed by restriction digestion using *Xba*I and *Hind*III enzymes. Selected colonies for each construct were confirmed by Eurofins sequencing. The Level 1 modules, *UB10::gORFcycp3;1:4xMyc:NOS* and *UB10::gORFcycp3;2:4xMyc:NOS*, in-planta *HYG* resisten cassette (pICSL11059) and linker Ele2 (pICH41744) were assembled into Level 2 acceptor backbone (pICSL4723) by the “dig-lig protocol” for the final Level 2 asembly using *Bbs*I (*Bpi*I) and T4 DNA ligase enzymes. Incubation of reaction in a thermocycler (PCR), bacterial transformation, screening of colonies (Kan^R^ LB plates) followed by restriction digestion (using *Hind*III & *Pst*I) and confirmation of colonies by Eurofins sequencing was done as aforementioned. Col-0 plants were transformed with each of the two construct using the flower dip method(*56*) and *Agrobacterium tumefaciens* strain *GV3101*. Positive transformants were selected on Hyg. In T1, Hyg-resistent plants segregating 1:4 were selected (one copy of the construct). In T2, homozygote Hyg-resistent plants were selected and used for further analysis.

#### SPT-driven overexpression of CYCP3;2 and IPT7 lines

*pSPT:gCYCP3;2:HA* construct was generated by In-Fusion cloning as follows: L0 gORF of *CYCP3;2* as constructed above for *UB10::CYCP3;2:Myc* construct was digested with a primer pair CYCP3;2-F (KpnI) /CYCP3;2-R (XhoI) (see Suppl. table 2) and confirmed by Eurofins sequencing. CYCP3;2 was then cloned into pre-digested linearized *pSPT–pCambia1305-3xHA* (4980bp) vector (gifted by Yuxiang Jiang from host lab)(*12*) by In-Fusion ligation reaction using 5x In-Fusion® HD enzyme, thus generating *pSPT:CYCP3;2:HA* construct. All constructs were confirmed by sequencing and transformed into *Agrobacterium tumefaciens* strain *GV3101* for plant transformation in Col-0 background. Positive T1 lines were selected on Basta^R^ MS plates and used for further analysis. *pSPT:IPT7* construct (for transformation into Col-0, *trm6,7,8*, and *ton1a*) was constructed similarly. Complete coding sequence (CDS) of *IPT7* gene (990bp) including the stop codon was amplified from Col-0 CDS DNA using gene specific primers (see Suppl. table 2). *IPT7* CDS sequence was digested with primers pair IPT7-F (KpnI)/IPT7-R (XhoI) and cloned into pre- digested linearized *pSPT–pCambia1305-3xHA* (4980bp) vector by In-Fusion ligation reaction using 5x In-Fusion® HD enzyme. The absence of mutations was verified by Eurofins sequencing and the construct was transformed into *Agrobacterium tumefaciens* strain *GV3101* for plant transformation. Positive T1 lines were selected on Basta^R^ MS plates and used for further analysis.

#### cycp3;1-1 and cycp3;2-1 CRISPR mutants

Single CRISPR Cas9 mutants for CYCP3;1 (*cycp3;1-1*) and CYCP3;2 (*cycp3;2-1*) were obtained as follow: Two guides were used for each gene (guide3: CTAGGAACGAGAGAATCAGC and guide1: GTATACCAAAGCCGGTCCAT for CYCP3;1; guide9: TGACCATCCAGTCATACCTA and guide6: GTACACTAAAGCCGGTCCTT for CYCP3;2) and included in specific forward primers to be cloned using the Golden Gate cloning technology. Each forward primer was used in combination with a universal reverse primer for PCR amplification using Phusion Polymerase Taq (30 cycles at 56°C for 10sec). Amplification products were separated on a 2.5% agarose gel and purified using gel filtration kit (Sigma). CYCP3;1 guide3 and CYCP3;2 guide9 were cloned into pICH47751 vector, while CYCP3;1 guide1 and CYCP3;2 guide6 were cloned into pICH47761 vector. All resulting constructs were transformed in *E. coli* competent cells (DH5α) and selected on LB medium (Carb^R^). After plasmid extraction using NucleoSpin^®^ Plasmid kit from the positive colonies and sequencing by Eurofins, the two guides for each gene were combined in a L2 reaction using the pICSL4723 destination vector, alongside the pICSL11015 (FastRed selection in plants), the CAS9 BCJJ358 vector, and the linker Ele4 pICH41780. The *E. coli* positive colonies were selected on Kan^R^ LB plates. L2 plasmids were used to transform Col-0 plants using the *Agrobacterium* infiltration methods. Individual T1 FastRED positive seeds were selected using a stereo fluorescent microscope (Leica M205FA). Genomic DNA was extracted by individual T1 and T2 adult plants using the isopropanol method. Each DNA was used as a template to amplify a region across the two guides: for *cycp3;1-1*, CYCP3;1_CRISPRseq_F and CYCP3;1_CRISPRseq_R (see Suppl. table 2) were used to amplify a region of 851bp, while for *cycp3;2-1*, CYCP3;2_CRISPRseq_F and CYCP3;2_CRISPRseq_R (see Suppl. table 2) were used to amplify a region of 723bp. The PCR products were partially run on a 1.8% agarose gel and the rest was cleaned up for sequencing using the respective CRISPRseq_F primers. Scrambled sequences at the guide positions onwards were considered edited and the corresponding plants offsprings were grown to obtain a second generation of edited plants. In T2s, FastRED negative seeds were selected to eliminate the Cas9, the genomic DNA was extracted from adult plants and amplified as above. Sequencing results revealed homozygosis for a point mutation for *cycp3;1-1* (line6-4) and a big deletion for *cycp3;2-1* (line B2-1) (see Suppl. Figure 3).

Homozygote *cycp3;2*-1 was crossed to *spt-12* mutant and the resulting double mutant was crossed to homozygote *cycp3;1-1* mutant to obtain a *spt cycp3;1 cycp3;2* triple mutant as well as a segregating *cycp3;1 cycp3;2* double mutant. Homozygosis for both CRISPR *CYCP3s* alleles was tested by genomic DNA extraction followed by sequencing, while the *spt-12* wildtype and T-DNA alleles were screened by PCR using the primer pairs spt-12 RP/spt-12 LB for wildtype and spt-12 RP/spt-12 LP for T-DNA (see Suppl. table 2).

#### *pCYCP3;1:GUS* and *pCYCP3;2:GUS* transcriptional fusion lines

A fragment of 2949bp upstream of the CYCP3;1 start codon was cloned to produce the GUS transcriptional fusion *pCYCP3;1:GUS* using the primers pair pCYCP3;1_FWD/ pCYCP3;1_REV (see Suppl. table 2), while a fragment of 2845bp upstream of the CYCP3;2 start codon was cloned to produce the GUS transcriptional fusion *pCYCP3;2:GUS* using the primers pair pCYCP3;2_F/ pCYCP3;2_R (see Suppl. table 2). The PCR products were cloned into the destination L0 vector pICH41295 using the Golden Gate cloning strategy and then transformed into *E. coli* (DH5α). Positive colonies were selected on LB plates supplemented with Spec. Plasmid DNA was extracted using NucleoSpin^®^ Plasmid kit, enzymatic digestions were performed to confirm the correct size of the inserts within the receiving plasmids, and sequencing by Eurofins. L1 reactions were performed to combine each promoter to the GUS reporter by using the pICH47742 backbone, the pICH75111 vector containing the GUS sequence and the pICH41421 NOS terminator. The two resulting constructs were transformed in *E. coli* (DH5α) and the positive colonies screened on Carb^R^ LB plates. Enzymatic digestions and sequencing were performed to check the size and the correct junction of the GUS reporter to the promoters. L2 reactions were then performed to clone the *pCYCP3;1:GUS* and *pCYCP3;2:GUS* constructs in to the destination vector pAGM4273 alongside the vector pICSL11024 for the resistance cassette (Kan^R^) for selection in plants. The resulting vectors were transformed into *Agrobacterium tumefaciens* strain *GV3101* before transformation into Col-0 plants. T1 with sinlge insertion and homozygote T2 transgenic lines were selected using resistance to KAN. Homozygous T2 lines were then crossed to the *spt-12* and *ind-2* mutants to obtain double homozygote lines (*spt-12 pCYCP3;1:GUS; ind-2 pCYCP3;1:GUS; spt-12 pCYCP3;2:GUS* and *ind-2 pCYCP3;2:GUS*). Also *pCYCP3;1:GUS* and *pCYCP3;2:GUS* were crossed to *35S::ARR1*Δ*DDK:GR* to generate several F1s to be used for further analysis.

#### Cloning strategy for co-expression analysis in *N. benthamiana*

Constructs of *35S::SPT-RFP* and *35S::NLS-RFP* were created as follow: the full-length coding sequences of *SPT* and *NLS* were digested with SifI (NEB) and cloned into the empty *pCambia1305–35S::RFP* vector respectively, which was pre-digested with DraIII (NEB). For the genomic *CYCP3;1* and *CYCP3;2* constructs (without STOP codon), Golden Gate cloning assembly was used as described above. The *CYCP3;1* promoter (2949bp upstream of the start codon) and *CYCP3;2* (2845bp upstream of the start codon) were cloned as described above. Genomic fragments of CYCP3;1 and CYCP3;2 transcript regions without the stop codon were amplified and cloned into pAGM1287. These entry clones were combined with CITRINE (Cit) in pAGM1301 and inserted into the L1 vector pICH47742 and finally into the destination vector pAGM4723 together with pICSL11059 that confers the Hygromycin resistance cassette. For generation of the constructs harbouring different G-BOX mutations of the promoters, p*CYCP3;1:CYCP3;1-CITRINE* and p*CYCP3;2:CYCP3;2-CITRINE* in pAGM4723 were used as templates respectively to introduce specific point mutations in the promoter sequence with mutagenesis primers (listed in Suppl. table 2). These binary constructs were introduced into *Agrobacterium tumefaciens GV3101* strain for infiltration in *N. benthamiana* leaves.

### Chromatin immunoprecipitation sequencing (ChIP-seq)

Young inflorescences were chopped off from 4 weeks old plants of *spt-12/SPT*::*SPT*–*sYFP* line(*12*), and immediately frozen in liquid nitrogen after collection as described previously(*12*). 3g of inflorescences were used for each biological replicate; experiment was performed with 3 biological replicates. Further ChIP assay was performed as described previously(*57*) (Andre’s). Immunoprecipitation (IP) was conducted using GFP-Trap® magnetic particles M-270 (ChromoTek). IP and input samples (n=6, 3 for each IP and input) were sequenced by Novogene Illumina Sequencing (PE150). Raw reads were processed and trimmed using fastp (v0.20.1) and aligned to the TAIR10 genome with Bowtie2 (v2.5.1)(*58*). The mapped data was sorted, indexed; duplicate reads were flagged using Samtools(*59*) (v1.9). Reads overlapping blacklisted genome regions, as defined by the Greenscreen Project(*60*), were removed using Bedtools(*61*) (v2.31.0). Peaks were called using MACS3 (v3.0.0a7)(*62*) with the parameters callpeak -p 0.1 -B --bdg -- keep-dup auto. The resulting peaks were further filtered based on different cutoffs and submitted to PAVIS(*63*) for annotation and visualization (https://manticore.niehs.nih.gov/pavis2/annotate). Peaks with a 0.001 false discovery rate (FDR) were assigned to gene models within 2kb upstream and 1.5kb downstream regions. Target genes were shortlisted if they appeared in all three replicates. ChIP-seq data was visualized using the Integrative Genomics Viewer(*64*). The peaks and the genome tracks were plotted using pyGenomeTracks(*65, 66*). The GO enrichment and visualisation was done using ShinyGO(*67*) v0.80 (bioinformatics.sdstate.edu/go) with FDR cutoff 0.05 against *Arabidopsis thaliana* TAIR10 assembly.

### ChIP-qPCR Assay

Following ChIP protocol as described above, the enrichment of CYCP3;1, CYCP3;2 and CYCD1;1 promoter regions (around the G-boxes cis-elements) wasquantified using qPCR. Enrichment values were normalized against *ACTIN7* gene. Primers used for the enrichment of each gene are listed in Suppl. table 2.

### RNA extraction and qRT-PCR

For qRT-PCR analysis, RNeasy Plant Mini Kit (Qiagen) was used to extract total RNA from either young inflorescences or 7 days old seedlings in triplicate (3 independent biological replicates). Reverse transcription of extracted RNA was done using M-MLV Reverse Transcriptase (Promega). At least 3 independent technical experiments were performed from each RNA sample using SYBR Green Master Mix (Promega) with Chromo4 Real-Time PCR Detection System (Bio-Rad). Target gene expression levels were normalised against *UBIQUITIN10*. Relative expression levels were quantified using the 2^-ΔΔCT^ method in Microsoft Excel (v.2311). The results from one representative experiment are plotted in figures. The gene specific primers used for the analysis are listed in Suppl. table 2.

### Scanning electron microscopy

Whole inflorescences were fixed overnight in FAA (3.7% formaldehyde, 5% glacial acetic acid, 50% ethanol) and dehydrated through an ethanol series (50% to 100%) as described previously(*3*). After complete dehydration, samples were critical point-dried using Leica EM CPD300. Gynoecia were dissected manually using stereomicroscope (Leica S9D) and mounted on stubs. Samples were sputtered coated using ACE-600 before examination using FEI Nova NanoSEM 450 emission scanning electron microscope. An acceleration voltage of 3 kV was used for imaging samples. The total number (n) of gynoecia imaged for each experiment are mentioned in the figures’ legends.

### GUS histochemical analysis

To analyse *pCYCP3;1:GUS* and *pCYCP3;2:GUS* lines expression in wildtype (Col-0) and mutant/overexpression (*spt-12*, *ind-2*, *35S::ARR1*Δ*DDK:GR*) backgrounds, whole inflorescences were chopped off plants and acetone pre-treatment and GUS-staining was performed as described previously(*3*). For expression analysis of *pCYCP3;1:GUS* (T2), *spt-12 pCYCP3;1:GUS* (F2), *ind- 2 pCYCP3;1:GUS* (F2) *and 35S::ARR1*Δ*DDK:GRxpCYCP3;1:GUS* (F1), whole inflorescences were stained for 4.5hrs. While, for visualization of *pCYCP3;2:GUS* (T2), *spt-12 pCYCP3;2:GUS* (F2), *ind-2 pCYCP3;2:GUS* (F2) *and cxpCYCP3;2:GUS* (F1), staining was performed overnight (24hrs). Samples were washed with sterile water after decanting GUS-solution and replaced with few dilutions of 70% ethanol until chlorophyll was completely removed. Gynoecia were manually dissected using stereomicroscope (Leica S9D) and mounted on glass slides in 8:3:1 chloralhydrate (Sigma-Aldrich) solution. Mounted samples were analysed by Zeiss Axio Imager Z2 light microscope using DIC prisms.

### Transient expression assay in tobacco leaves

The transient co-expression assays were performed on 4-wk-old *N. benthamiana* plants infiltrated with *Agrobacterium tumefaciens* strains carrying the respective binary expression plasmids. *A. tumefaciens* suspensions were prepared in infiltration buffer (10 mM MES, 10 mM MgCl_2_, and 150 μM acetosyringone, pH 5.6) and were adjusted to appropriate OD_600_. *Agrobacterium* strain harbouring P19 was also co-infiltrated to enhance gene expression. After 48 h of infiltration, the infiltrated tobacco leaves were observed under the confocal laser scanning microscope.

### Confocal microscopy

For confocal imaging of tobacco leaves, a Zeiss LSM 880 confocal scanning microscope was used with the following fluorescence excitation–emission settings to visualize: CITRINE (*Cit*) excitation 514 nm, emission 530 nm; RFP excitation 550 nm, emission 580 nm. Pictures were taken with 20× or 40× water/oil immersion objectives. Samples within one experiment were imaged with identical settings. For image analyses, ImageJ and Zeiss Zen 2011 (v3.4) image analyses software were used.

### Statistical analysis

Relative expression levels were compared using Student’s unpaired *t*-test. *p* value <0.05 was considered significant. For comparison of phenotypic classes, 2x2 contingency tables were generated, followed by Fisher’s exact Chi^2 test. Two-tailed p values <0.0001 were considered extremely statistically significant. Experimental data was obtained by counting the number of phenotypes, while their percentage is plotted in the graphs. For style length and width comparison, on-way ANOVA (analysis of variance) followed by Tukey’s Honestly significance difference (HSD) was used for pairwise comparisons. Tukey’s HSD p values <0.001 were considered extremely statistically significant. Photoshop® was used to assemble the Figures.

## Acknowledgments

We thank Dr David Bouchez and Dr Martine Pastuglia (INRAE, FR) for kindly providing the *trm6,7,8* triple mutant; Dr Silvia Costa (JIC, UK) for donating the *ton1a* and *pok1,2* mutants; Prof. Sabrina Sabatini (Sapienza University, IT) for providing the *35S::ARR1*Δ*DDK:GR* line. We also thank Dr Yuxiang Jiang, Dr Benguo Gu and Dr Anna Schulten (JIC) for technical help with the ChIP extraction protocol and the JIC bioinformatic platform.

## Funding

This research was funded by the Royal Society University Research Fellowship URF\R\231023 (LM), the Royal Society Enhanced Research Expenses RF\ERE\210323 (LM), the Royal Society Research Fellows Enhancement Awards RGF\EA\181077 (LM), and the Institute Strategic Programme grant (BB/X01102X/1) to the John Innes Centre from the Biotechnology and Biological Sciences Research Council. This work has benefited from the equipment and framework of the COMP-R Initiative, funded by the ‘Departments of Excellence’ program of the Italian Ministry for University and Research (MUR, 2023-2027).

**Author contributions**

L.M., conceptualised the project; L.M. and I.J. designed the experimental research; I.J., performed most of the experimental work with help from S.W.H.K, J.C. and L.M. All authors analysed the data. L.M. prepared the figures and wrote the manuscript. All authors commented and edited the manuscript.

**Competing interests**

The authors declare no competing interests.

**Data and materials availability**

The *spt-12/SPT*::*SPT*–*sYFP* raw ChIP-seq datesets are available via EBI/NCBI website under study accession number PRJEB80813 (https://www.ncbi.nlm.nih.gov/bioproject/?term=PRJEB80813). All processed data are contained in the manuscript or in the Supplementary information. Biological material and data from this study will be available upon request and with no restrictions.

## Supplementary Material

**Fig. S1.**
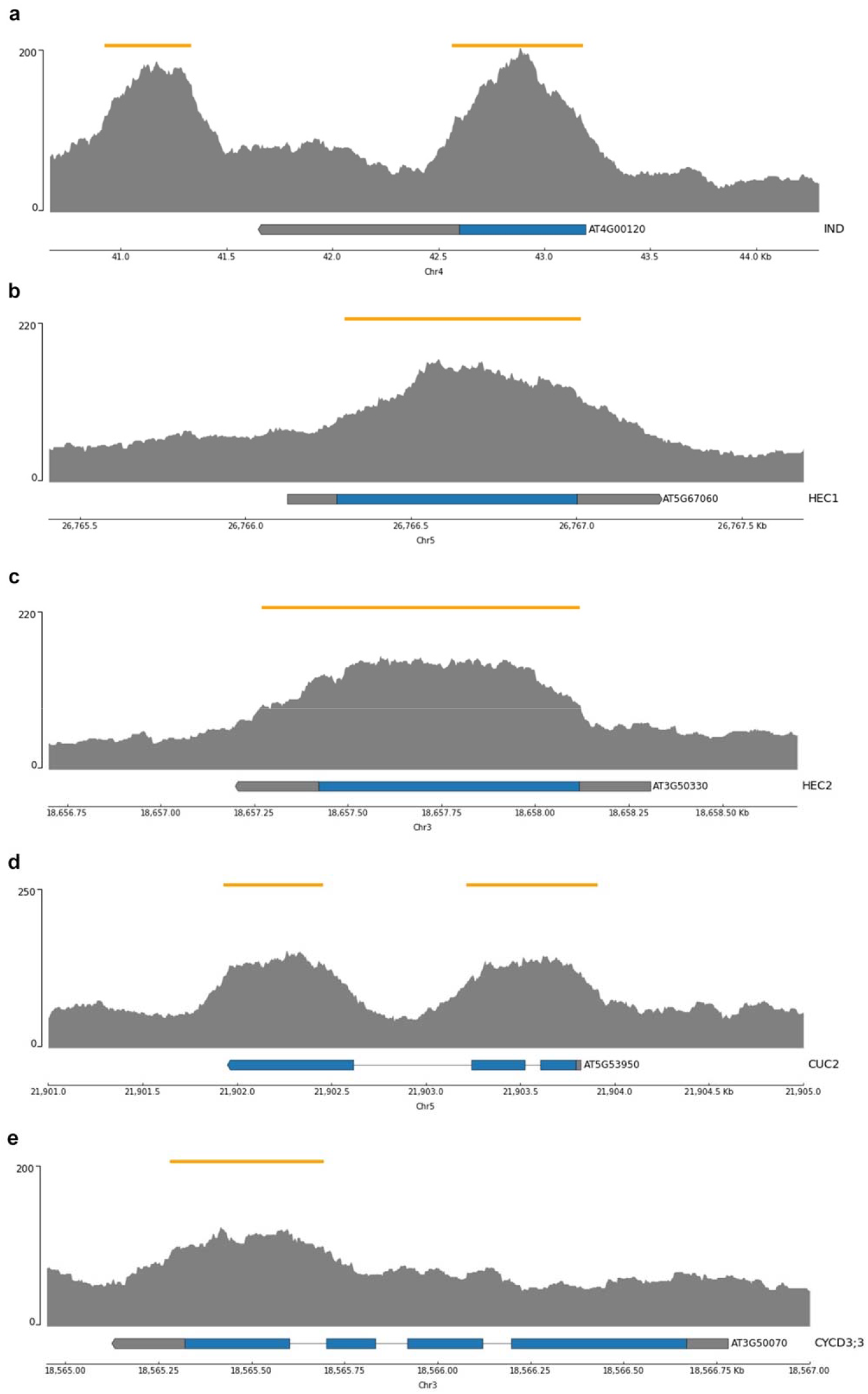
Presumptive direct targets of SPT from ChIP-seq experiments. (A-E) Representative raw Chromatin Immunoprecipitation sequencing (ChIP-seq) peaks of *IND* (A), *HEC1* (B), *HEC2* (C), *CUC2* (D) and *CYCD3;3* (E). n=3 biological replicates; peaks of one representative replicate are shown. Yellow bars on top represent peaks position on chromosome. Blue bars on bottom show exons, and grey lines show introns.

**Fig. S2.**
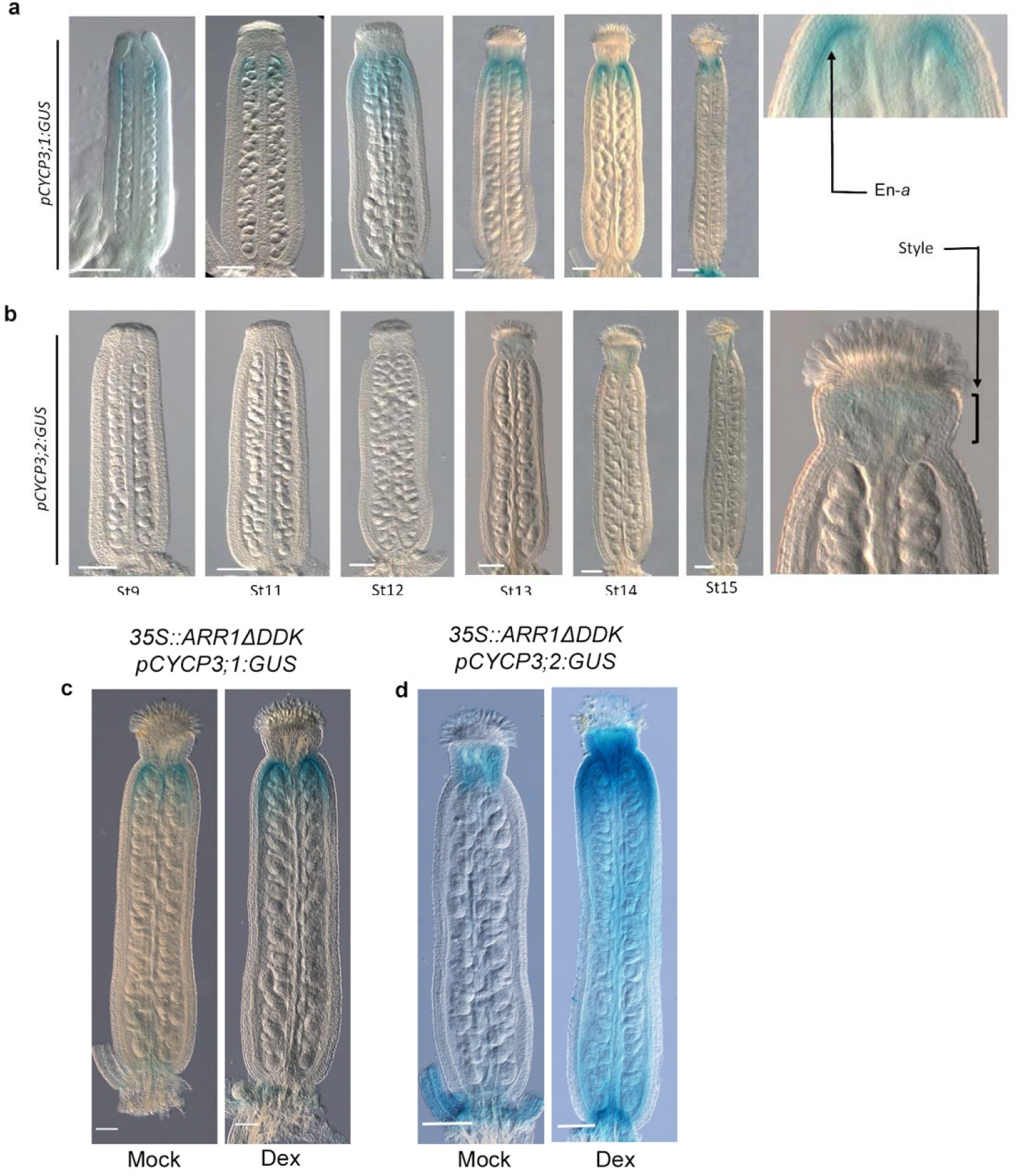
Expression of CYCP3s during gynoecium developmental stages and in *35S::ARR1*Δ*DDK:GR* background. (A, B) Light microscope images of GUS-stained gynoecia (stages 9-15) showing expression of *pCYCP3;1:GUS* (a) and *pCYCP3;2:GUS* (b) in wildtype (Col-0) background. Notice the expression of *pCYCP3;1:GUS* in endocarp-*a* (En-*a*) throughout developmental stages, and a very weak signal of *pCYCP3;2:GUS* in style of stage-15 gynoecia. Scale bars are 100 µm. 50 gynoecia were analysed for each transgenic line. n=3 biological repetitions. (C, D) Light microscope images of GUS-stained mock- (left) and DEX- (right) treated (10 µM) gynoecia of F1 *pCYCP3;1:GUS x 35S::ARR1*Δ*DDK:GR* and *pCYCP3;2:GUS x 35S::ARR1*Δ*DDK:GR.* 20 to 40 gynoecia were analysed for each F1 cross. n=3 biological repetitions. Scale bars are 200 µm.

**Fig. S3.**
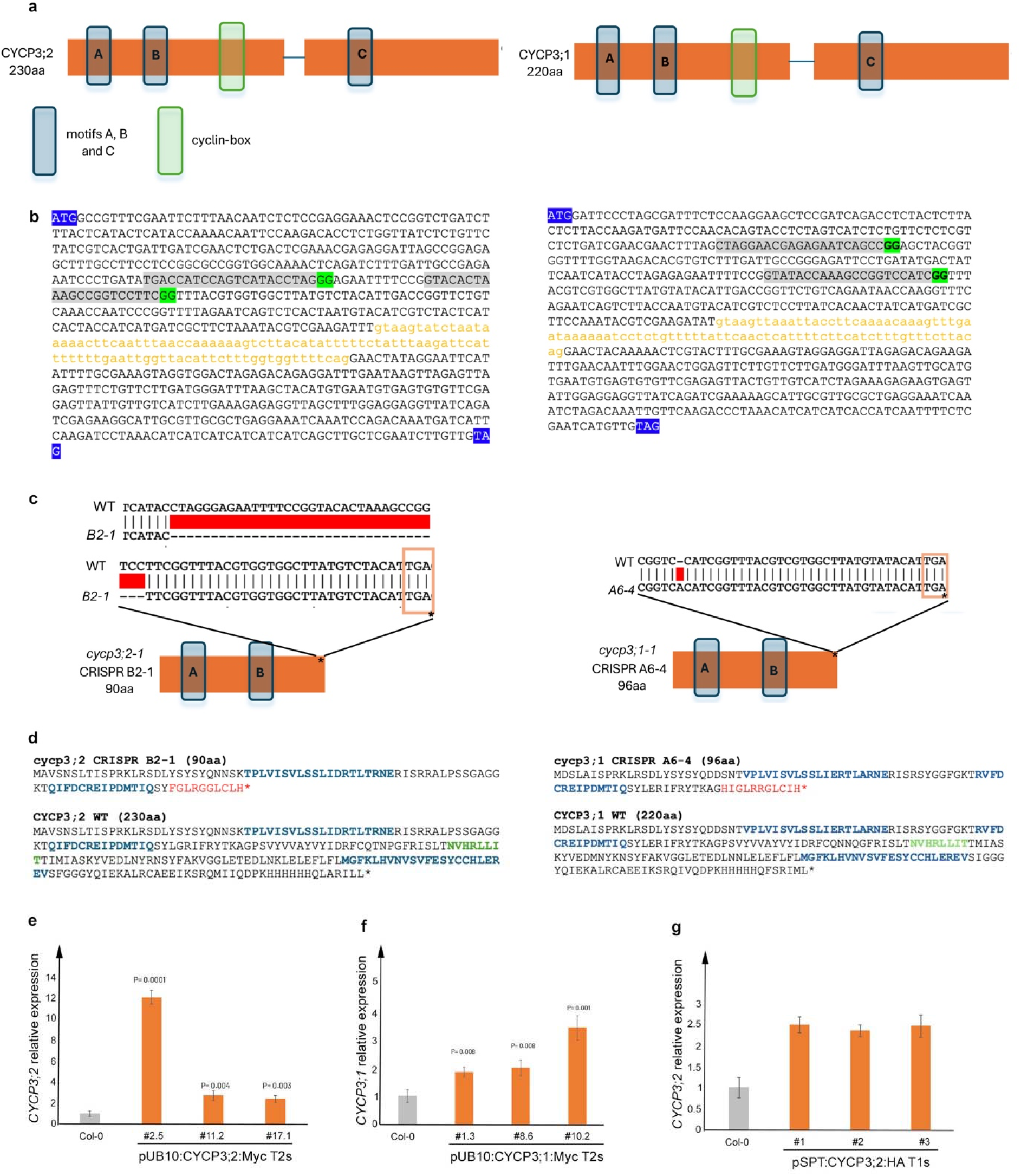
Cloning strategy for *CYCP3s* CRISPR mutants and expression levels of *CYCP3s* overexpressing lines. (A) Schematic representation of CYCP3;2 (left) and CYCP3;1 (right) protein structures showing exons (orange boxes), intron (blue line), the A,B,C motifs (blue boxes) and the cyclin-box protein domain (green box). (B) Genomic nucleotide protein sequence of CYCP3;2 (left) and CYCP3;1 (right) showing the position of the two guides (highlighted in grey) used to obtain the single *cycp3;2-1* and *cycp3;1-1* CRISPR mutants. The PAM recognition site (GG) is highlighted in green, the start and stop codons are highlighted in blue, exons are represented by black font while the intron is in orange font. (C) Schematic representation of CYCP3;2 (left) and CYCP3;1 (right) CRISPR products obtained by DNA sequencing of T2 plants. The scheme includes the alignment of WT and mutated sequences and shows the position of the big deletion recovered in *cycp3;2-1* (line #B2-1) and of a single-nt (nucleotide) insertion in *cycp3;1-1* in the first exon after the A and B motifs and before the cyclin-box domain. (D) Prediction of the amino acid sequence of *cycp3;2-1*(90 aa, left) and *cycp3;1-1* (96 aa, right) generated by CRISPR-Cas9 compared to the respective wild-type sequences (230 aa for CYCP3;2 and 220 aa for CYCP3;1), showing the position of the A and B motifs (blue font) and the shifted open reading frame (red font) generated by the mutations. Asterisks represent STOP codons. (E, F) Bar charts of qRT-PCR quantification showing relative expression of *CYCP3;2* (e) and *CYCP3;1* (f) in wildtype (Col-0) background compared to three independent overexpressing homozygous lines (T2s) of *UB10::CYCP3;2:Myc* (#2.5, #11.2 and #17.1) and *UB10::CYCP3;1:Myc* (#1.3, #8.6 and #10.2). Expression levels were normalised against *UBIQUITIN10*. n= 3 biological replicates. Error bars represent SD. Statistical significance, *p* values (Student’s *t-*test) are shown on each bar. (G) Bar charts of qRT-PCR quantification showing relative expression levels of *CYCP3;2* in Col-0 compared to three independent transgenic lines (T1s) of *pSPT:CYCP3;2:HA*. Expressions were normalised against *UBIQUITIN10*. The experiment was performed once on three independent transgenic lines, with four technical repeats. Values shown are meanslJ±lJSEM.

**Data Table S1.** (separate excel file) Analysis of the *spt-12/SPT*::*SPT*–*sYFP* ChIP-seq experiments conducted in triplicates using inflorescent material.

**Data Table S2.**
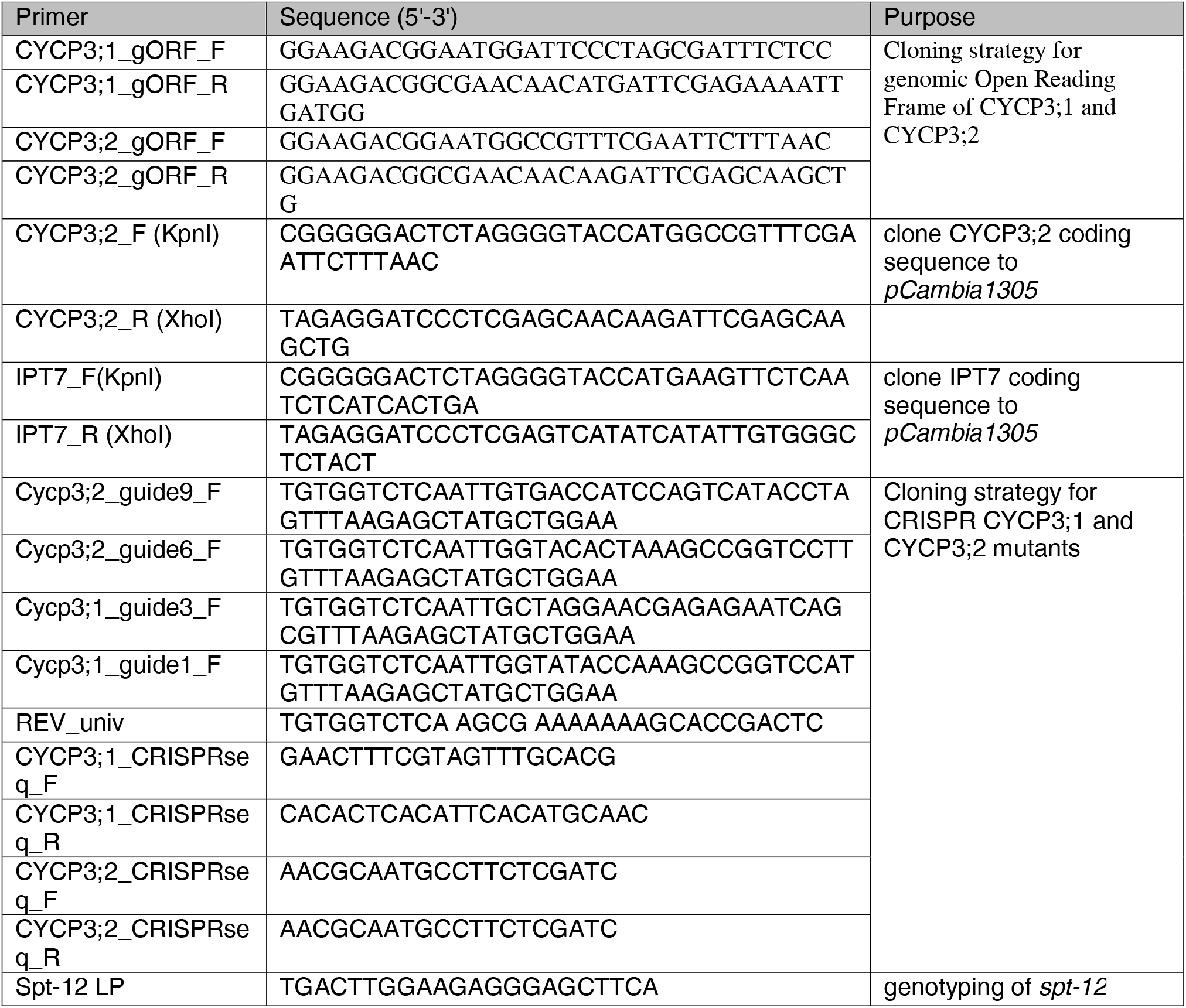
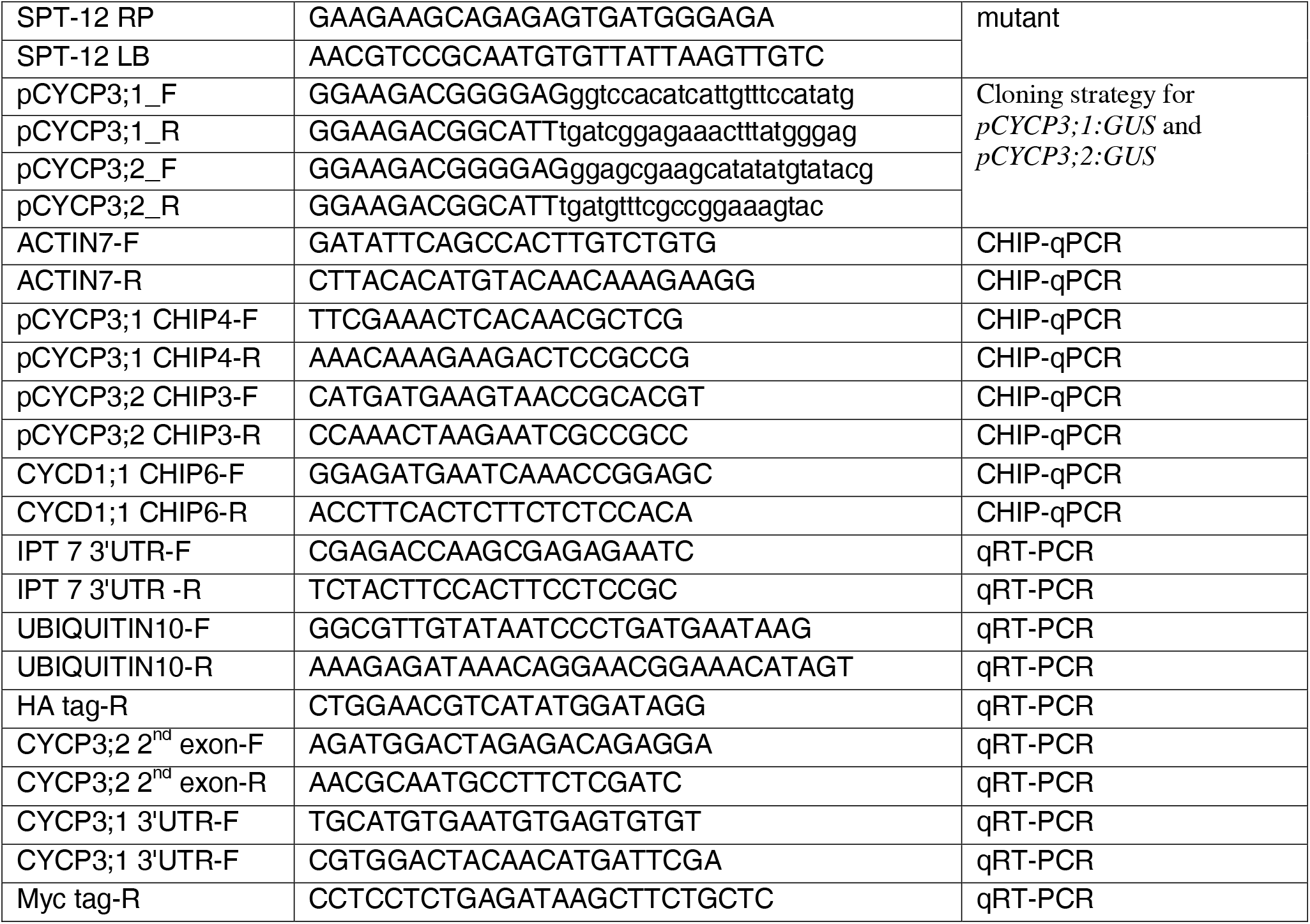
(separate excel file) List of primers used in the study

